# Combined epigenetic and metabolic inhibition blocks platinum-induced ovarian cancer stem cell enrichment

**DOI:** 10.1101/2021.03.18.435878

**Authors:** Riddhi Sood, Shruthi Sriramkumar, Vaishnavi Muralikrishnan, Sikai Xiao, Weini Wang, Christiane Hassel, Kenneth P. Nephew, Heather M. O’Hagan

## Abstract

High grade serous ovarian cancer (HGSOC) is the most common and aggressive type of ovarian cancer. Platinum resistance is a common occurrence in HGSOC and a main cause of tumor relapse resulting in high patient mortality rates. Recurrent OC is enriched in aldehyde dehydrogenase (ALDH)+ ovarian cancer stem cells (OCSCs), which are resistant to platinum agents. We demonstrated that acute platinum treatment induced a DNA damage-dependent decrease in BRCA1 levels. In a parallel response associated with G2/M arrest, platinum treatment also induced an increase in expression of *NAMPT*, the rate limiting regulator of NAD^+^ production from the salvage pathway, and levels of NAD^+^, the cofactor required for ALDH activity. Concurrent inhibition of DNA methyltransferases (DNMTs) and NAMPT synergistically abrogated the platinum-induced increase in OCSCs. Combining pharmacological inhibitors of DNMT and NAMPT with carboplatin reduced tumorigenesis and OCSC percentage *in vivo*. We conclude that both epigenetic and metabolic alterations lead to platinum induced OCSC enrichment, providing preclinical evidence that in the neoadjuvant setting, combining DNMT and NAMPT inhibitors with platinum has the potential to reduce OC recurrence and avert the development of platinum resistance.

## Introduction

While ovarian cancer (OC) is initially highly responsive to chemotherapy, recurrence is common, and recurrent OC is chemotherapy resistant and fatal (1). A subpopulation of cells called ovarian cancer stem cells (OCSCs) preferentially survive after platinum-based chemotherapy (2), are enriched in recurrent tumors (3) and are at least partially responsible for chemotherapy resistance (4). Several markers have been used to identify OCSCs, including the activity of aldehyde dehydrogenase (ALDH) enzymes. ALDH1 is overexpressed in OCSCs and correlates with worse survival and platinum resistance (2, 5). ALDH+ cells have tumor initiating capacity, form spheroids in non-adherent conditions, and express stemness genes (2, 6), all features of CSCs.

Platinum-based chemotherapeutic drugs damage DNA by forming platinum-DNA adducts (7), which activate the DNA damage response (DDR). The tumor suppressor breast cancer 1 (BRCA1) plays an important role in regulating the DDR through interaction with proteins required for cell cycle regulation, tumor suppression, and DNA repair (8-11). While about 40% of women with a family history of OC have *BRCA1/2* mutation or promoter DNA hypermethylation making them more sensitive to chemotherapy, the majority of OCs have wildtype *BRCA1* (12). Therefore, a better understanding of the role of wildtype BRCA1 in response to platinum agents is essential.

The metabolite nicotinamide adenine dinucleotide (NAD^+^) plays a key role in several metabolic pathways (13). Furthermore, NAD^+^ is a co-factor for ALDH enzymes (14, 15). OCs with reduced expression of *BRCA1* have increased levels of both NAD^+^ and expression of the rate limiting regulator of NAD^+^ synthesis salvage pathway, nicotinamide phosphoribosyltransferase (NAMPT) (16). NAMPT promotes platinum-induced senescence-associated OCSCs (17), further suggesting a connection between NAD^+^ and BRCA1 in the development of OC chemoresistance.

With the known involvement of OCSCs in chemoresistance and tumor recurrence, we sought to mechanistically study how platinum treatment induces OCSC enrichment and develop strategies to combat this enrichment. We demonstrate that decreased expression of BRCA1 and altered NAD+ levels function in parallel to drive platinum-induced OCSC enrichment. Cisplatin treatment resulted in a DDR-dependent decrease in *BRCA1* expression as well as a G2/M cell cycle arrest-related increase in *NAMPT* expression and subsequent increase in cellular NAD^+^ levels. Importantly, combined treatment with DNA methyltransferase (DNMT) and NAMPT inhibitors synergistically abrogated cisplatin-induced OCSC enrichment. Our in vitro and in vivo findings support combining epigenetic and metabolic inhibitors in the neoadjuvant setting to reduce platinum-induced enrichment of OCSCs and avert development of platinum resistance.

## Materials and methods

### Cell lines, culture conditions, and reagents

High grade serous OC (HGSOC) cell lines OVCAR5 (RRID:CVCL_1628), OVCAR3 (RRID:CVCL_0465), COV362 (RRID:CVCL_2420), OVSAHO (RRID:CVCL_3114) and PEO1 (RRID:CVCL_2686) were obtained from the Nephew lab, maintained using standard conditions and passaged for less than 15 passages (18-20). For most of the in vitro experiments, OHSAHO and OVCAR5 were used to represent a range (4.00 - 12.00 μM) of cisplatin sensitivity (20). All cell lines were tested for mycoplasma in 2017 (ATCC, 30-1012K) and authenticated by ATCC in 2018. Cisplatin (EMD Millipore, 232120) stock solutions was made in 154 mM NaCl at 1.67 mM. Cells were treated with cell line specific IC_50_ dose of cisplatin (OVSAHO, 4.00 μM; OVCAR5, 12.00 μM; PEO1, 12.84 μM; COV362, 13.57 μM; OVCAR3, 15.00 μM) (20). Cells were treated with CDK1i (9 μM for 16h; Sigma-Aldrich, SML0569) or decitabine (DAC, 100 nM for 48h; Sigma, A3656). Media containing fresh DAC was changed every 24h. Cisplatin was added during the last 16h of DAC treatment. Cells were treated with NAMPTi (50 nM for 6h; Sigma-Aldrich, SML1348). For cisplatin and NAMPTi dual treatment, cells were treated with cisplatin as above and NAMPTi was added 10h later. For low dose NAMPTi and DAC combination treatment with cisplatin, cells were treated with DAC (10 nM or 20 nM for 48h), cisplatin was added in the last 16h and NAMPTi (12.5 nM) was added during the last 6h of the DAC treatment. Cells were treated with ATM inhibitor KU-55933 (Sigma, MO #SML1109; 15 μM) for 16h in combination with cisplatin.

### ALDEFLUOR assay and flow cytometry

To measure ALDH activity, the ALDEFLUOR assay (Stem Cell Technologies, 01700) was used consisting of 1 million cells/1 mL ALDEFLUOR assay buffer and bodipyaminoacetaldehyde (BAAA) substrate +/- ALDH inhibitor diethylamino benzaldehyde (DEAB; 5 μL, 1.5 mM). Cells were incubated for 30-40 min at 37 °C, centrifuged and resuspended in ALDEFLUOR assay buffer.

Flow cytometry analysis was performed on a LSRII flow cytometer (BD Biosciences) at IU Flow Cytometry Core Facility. ALDH activity was measured using 488 nm excitation and the signal was detected using the 530/30 filter. For each experiment, 10,000 events were analyzed. ALDH+ percentage gate was determined by sample specific negative control (DEAB) ALDH+ gate. Further data analysis was done in FlowJo software (Becton, Dickinson & Company, RRID:SCR_008520).

### Cell cycle analysis

In order to analyze cell cycle in combination with the ALDEFLUOR assay, which requires live cells, nuclear ID red DNA stain (Enzo Life Sciences, ENZ-52406) was used. Cells were suspended in ALDEFLUOR reagent, incubated for 30 min at 37 °C, followed by incubation for 30 min in 1:250 dilution of Nuclear ID red stain in PBS at 37 °C and analyzed by flow cytometry. Nuclear ID red was excited at 561 nm and detected using the 670/30 filter.

### Quantitative RT-PCR (qRT-PCR)

Total RNA isolation and cDNA synthesis was performed as described previously (18). C_q_ values for genes of interest were normalized to housekeeping genes (*PPIA, β-Actin or RhoA*) using the deltaCq method. See Supplementary Table S1 for primer sequences.

### DNA extraction, bisulfite conversion, qMSP and bisulfite sequencing

DNA extraction, bisulfite treatment, and qMSP were performed as described previously (21). For bisulfite sequencing, bisulfite converted DNA was amplified and the PCR product was cloned into One Shot TOP10 Chemically Competent *E. coli* using the TOPO™ TA Cloning™ Kit (ThermoFisher, 451641). Plasmid DNA from bacterial colonies was extracted using Zyppy Plasmid Miniprep Kit (Zymo research, D4020) and sequenced by Sanger sequencing. Sequence peaks were analyzed for good quality in 4peaks software and DNA methylation maps were generated through BioAnalyzer (22). See Supplementary Table S1 for primer sequences.

### Western blot analysis

Cell pellets or pieces of xenografts were lysed in 4% SDS buffer using a QIAshredder (Qiagen, 79654). See Supplementary Table S1 for antibodies used. Band density was measured by ImageJ software (NIH, RRID:SCR_003070) and normalized to laminB, β- actin or vimentin.

### Spheroid formation assay

1.5 × 10^4^ cells pre-treated with cisplatin (6 μM for 3h), NAMPTi (50 nM for 6h), and/or DAC (100 nM for 48h) were plated in a 24-well low attachment plate (Corning, 3473) containing stem cell media (23) for 14 days. On day 14, images were taken using an EVOS FL Auto microscope (Life Technologies). To measure cell viability, Abcam ab176748 reagent, which measures cell viability be intracellular esterase activity, was added directly to each spheroid well for 1h. Viability (Ex/Em: 405/460 nm) was measured using a SynergyH1 plate reader (BioTek).

### NAD^+^/NADH ratio

NAD^+^/NADH ratio was calculated using NAD^+^/NADH quantification colorimetric kit (BioVision, K337-100) according to the manufacturer’s instructions.

### Transfection

Cells were transfected using Turbofect (ThermoFisher Scientific, R0532) 48h prior to treatment. pBABEpuro HA-BRCA1 was a gift from Stephen Elledge (Addgene plasmid # 14999, RRID:Addgene_14999) (24). pBABE-puro was a gift from Jay Morgenstern and Hartmut Land (Addgene plasmid # 1764, RRID:Addgene_1764) (25).

### Viral shRNA knockdown

BRCA1 knockdown was performed using shRNA1 (Sigma, TRCN0000244986) and shRNA2 (Sigma, TRCN0000244984) and empty vector (EV) TRC2 (Sigma, SHC201) followed by puromycin selection as previously described (18).

### Mouse Xenografts

All mouse experiments were approved by the Indiana University Bloomington Institutional Animal Care and Use Committee in accordance with the Association for Assessment and Accreditation of Laboratory Animal Care International. OVCAR3 cells (2 million) were injected s.c. into the flanks of NRG mice (NOD.Cg- *Rag1*^*tm1Mom*^ *Il2rg*^*tm1Wjl*^/SzJ; Jackson Laboratories; RRID:IMSR_JAX:014568). Once tumors were >100 mm^3^, mice were randomized and treated with vehicle, carboplatin (25 mg/kg weeks 1-2 and 50 mg/kg weeks 3-5; i.p. once weekly on day 3), carboplatin + DAC (0.1 mg/kg, i.p. once daily, 5 days per week), carboplatin + STF118804 (NAMPTi, 6.25 mg/kg s.c. twice daily, 7 days per week), or carboplatin + DAC + NAMPTi for 5 weeks. DAC was prepared in sterile PBS and STF118804 was prepared in 5% (v/v) DMSO and 20% (w/v) 2-hydroxypropyl-gamma-cyclodextrin (vehicle). Each group had 4-5 mice. At the end of the study, tumors were dissociated into single cells using a Tumor Dissociation Kit and a gentleMACS dissociator (Miltenyi Biotec) and used for ALDEFLUOR assays.

### Statistical methods

All experiments were performed in at least three biological replicates. When two groups were compared, statistical comparison was performed by Student’s t-test. One-way ANOVA followed by Tukey post hoc test was used to compare multiple groups using Graphpad Prism (RRID:SCR_002798).

## Results

### Cisplatin treatment enriches for ALDH+ cells

Advanced stage OC patients frequently have OCSC-enriched residual tumor cells following chemotherapy. To determine whether OCSCs are enriched by platinum chemotherapy, we treated HGSOC cell lines with corresponding IC_50_ doses of cisplatin (20) and analyzed the percentage of ALDH+ (%ALDH+) cells (2). In OVCAR5, the %ALDH+ cells significantly increased after 8h and 16h cisplatin treatment (Fig. 1A, Supplementary Fig. S1A). Similarly, 16h cisplatin treatment increased the %ALDH+ cells in OVSAHO (26), which have a homozygous deletion of *BRCA2*, OVCAR3 (wild-type *BRCA1/2*) and *BRCA2* mutant PEO1 cells (19) (Fig. 1A, Supplementary Fig. S1B).

**Fig 1.**
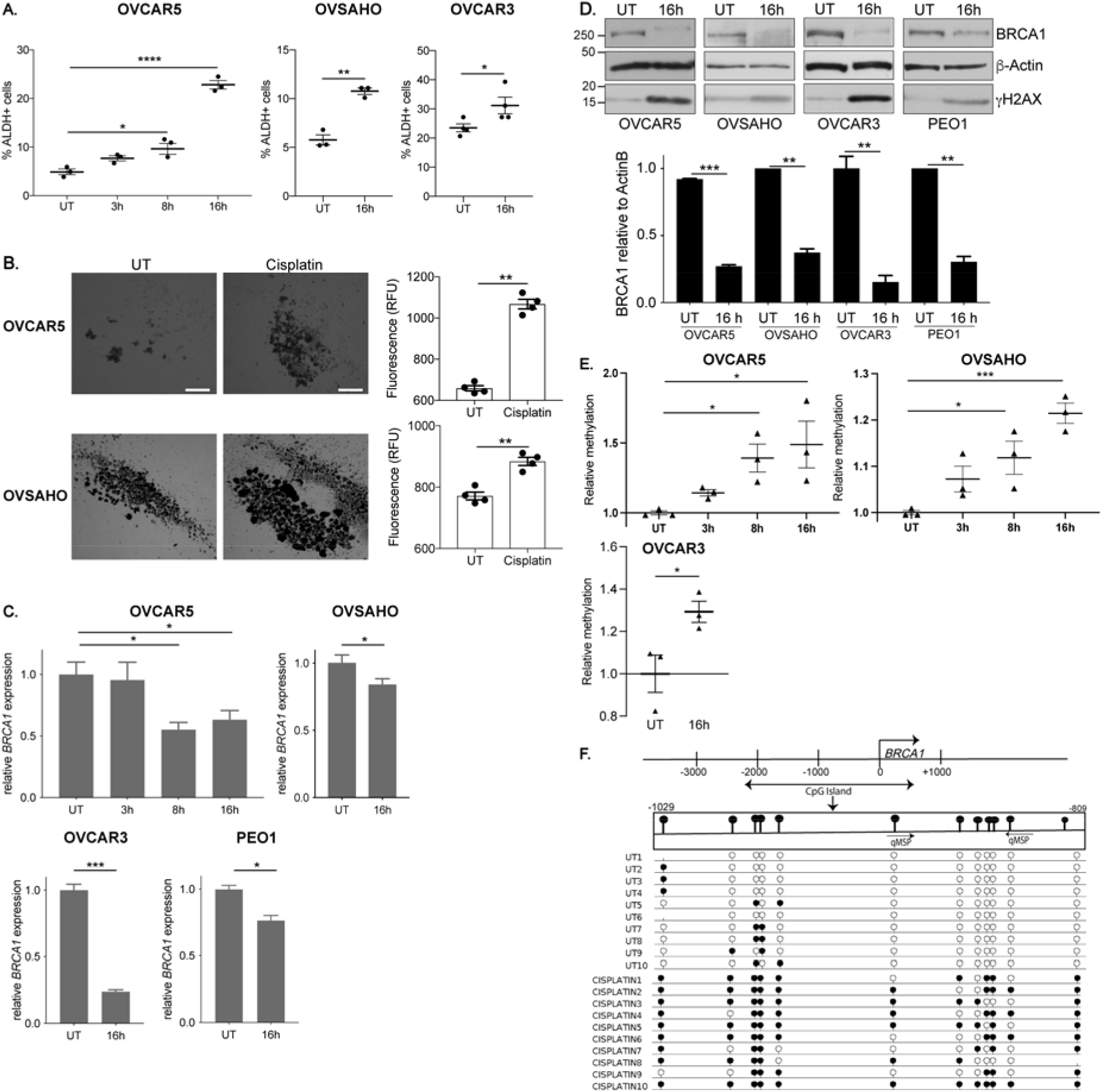
Cisplatin treatment enriches for ALDH+ cells. A) %ALDH+ cells after mock (UT) or cisplatin (IC50 dose, OVCAR5: 12 μM, OVSAHO: 4 μM, OVCAR3: 15 μM) treatment for the indicated time points. N=3. **B)** Images of spheroids after mock (UT) or cisplatin (3 h, ½ IC50, OVCAR5: 6 μM, OVSAHO: 2 μM) pretreatment in indicated cell lines. Scale bar = 500 μm. Graph depicts fluorescence intensity (RFU) of CytoCalcein Violet 450 stain. N=4. **C)** Relative *BRCA1* RNA expression in indicated cell lines after treatment as in A. N=3. **D)** Western blot and relative densitometry of whole cell lysates after treatment as in A. N=3. **E)** Quantitative MSP for *BRCA1* promoter DNA methylation after treatment as in A. N=3. **F)** The location and methylation status of CpG sites in promoter region of *BRCA1* gene. Location of qMSP primers are indicated. The arrow at the transcription start site (TSS) indicates transcription direction. Individual CpG dinucleotides are shown as circles with closed circles: methylation and open circles: unmethylated. Sequencing of ten individual clones each from bisulfite-converted DNA from mock (UT) or cisplatin (16h, IC50, 12 μM) cells. For all panels, graphs indicate mean +/- SEM, \**P*<0.05, \*\**P*<0.001, \*\*\**P*<0.0001, \*\*\*\**P*<0.00001.

However, in *BRCA1* mutant COV362 cells (27), no increase in %ALDH+ cells was observed after cisplatin treatment (Supplementary Fig. S1B). In anchorage-independent conditions, cisplatin pretreated cells were more spheroid-like (Fig. 1B) and an increased number of viable cells was observed compared to untreated cells (Fig. 1B), confirming that the increased %ALDH+ cells was associated with a stemness phenotype.

Isotypes of ALDH1A are linked to stemness of OC cells (28). We hypothesized that the observed cisplatin-induced enrichment of ALDH+ cells was due to altered expression of *ALDH*. However, no change in *ALDH1A1/A2/A3* isoform expression or ALDH1 protein levels was observed in cisplatin treated OVCAR5 cells (Supplementary Fig. S1C, D). In OVSAHO cells, no change in *ALDH1A1* expression, the major ALDH1 isoform, was observed after cisplatin treatment, although *ALDH1A2* and *ALDH1A3* isoforms significantly decreased and increased, respectively (Supplementary Fig. S1C).

As BRCA1 levels have been linked to an interstrand crosslink (ICL)-dependent increase in stemness (29), we assayed *BRCA1* expression after cisplatin treatment and observed significantly decreased *BRCA1* expression and protein levels in OVCAR5, OVSAHO, OVCAR3 and PEO1 cells (Fig. 1C,D). To determine if platinum resistant cells responded similarly to acute platinum treatment, we generated platinum resistant OVCAR5 cells by repeatedly exposing parental cells to the IC70 cisplatin dose. The resistant cells had a higher baseline %ALDH+ cells than the parental cells, which increased further with cisplatin treatment (Supplementary Fig. S1E). BRCA1 levels were higher in resistant cells than parental cells at baseline but decreased after cisplatin treatment at both the RNA and protein level (Supplementary Fig. S1F, G). Altogether, these data suggest that acute cisplatin treatment enriched for ALDH+ cells with stemness properties and decreased *BRCA1* levels.

### Cisplatin-induced decrease in *BRCA1* levels is associated with *BRCA1* promoter DNA hypermethylation

Various mechanisms have been reported to regulate *BRCA1* expression, including promoter DNA hypermethylation-associated gene silencing (30). Based on qMSP analysis, *BRCA1* promoter DNA methylation was increased significantly after 3h, 8h and 16h cisplatin treatment in OVCAR5 and OVSAHO cells and after 16h cisplatin treatment in OVCAR3 cells (Fig. 1E). Bisulfite sequencing of the *BRCA1* promoter region confirmed the increase in methylated CpGs after 16h cisplatin treatment compared to untreated (Fig. 1F, Supplementary Fig. S1H). Altogether this data demonstrates that the cisplatin-induced decrease in *BRCA1* expression is associated with promoter DNA hypermethylation.

### The cisplatin-induced decrease in BRCA1 is necessary for the associated increase in %ALDH+ cells

To determine if decreased *BRCA1* expression is sufficient to increase the %ALDH+ cells, BRCA1 was stably knocked down (KD) using shRNA. BRCA1 KD reduced *BRCA1* RNA and protein expression to levels similar to empty vector (EV) cells treated with cisplatin (Fig. 2A,B). BRCA1 shRNA1 KD cells had similar baseline %ALDH+ cells compared to EV (Fig. 2C). The slight increase in %ALDH+ cells in BRCA1 shRNA2 KD compared to EV cells was significantly less than the cisplatin-induced increase in EV cells.

**Fig 2.**
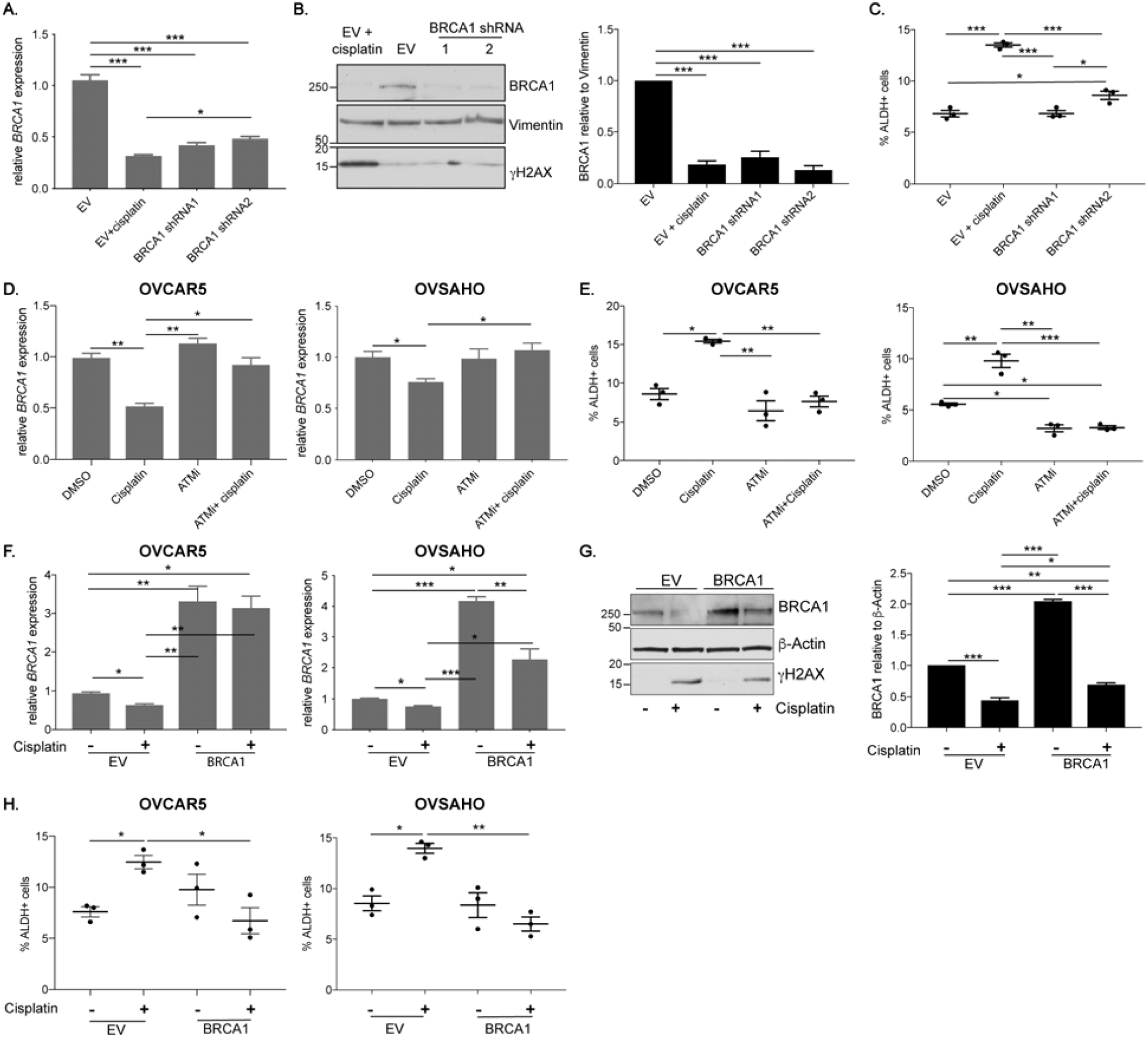
The cisplatin-induced decrease in BRCA1 is required for the associated increase in %ALDH+ cells. **A)** Relative *BRCA1* RNA expression in OVCAR5 cells after stable lentiviral infection with mock empty vector (EV) or BRCA1 shRNA1/2. N=3. **B)** Western blot and relative densitometry of OVCAR5 whole cell lysates after EV knockdown with or without cisplatin treatment (16h, IC50, 12 μM) or BRCA1 shRNA1/2 knockdown. N=3. **C)** Percentage of ALDH+ cells after stable EV knockdown with or without cisplatin treatment (16h, IC50, 12 μM) or BRCA1 shRNA1/2 knockdown. N=3. **D)** Relative *BRCA1* RNA expression after mock (DMSO), cisplatin treatment (16h, IC50, OVCAR5: 12 μM, OVSAHO: 4 μM), ATMi (KU-55933, 16h, 152 μM) or ATMi + cisplatin. N=3. **E)** Percentage of ALDH+ cells cells treated as in D. N=3. **F)** Relative *BRCA1* RNA expression in cells transfected with EV or CpGi-null BRCA1 plasmid and treated with or without cisplatin as in D. N=3. **G)** Western blot and relative densitometry of OVCAR5 whole cell lysates transfected and treated as in F. N=3. **H)** Percentage of ALDH+ cells after transfection and treatment as in F. N=3. For all panels, graphs indicate mean +/- SEM, \**P*<0.05, \*\**P*<0.001, \*\*\**P*<0.0001.

Because of the limited effect of decreasing BRCA1 levels on the %ALDH+ cells, we hypothesized that cisplatin DNA damage is important for the increase in %ALDH+ cells. Ataxia telangiectasia mutated (ATM), one of the first proteins to be recruited to DNA damage sites (31), is responsible for phosphorylation of downstream targets like H2AX and activation of downstream DNA repair pathways (32). Inhibiting ATM (ATMi) (33) reduced cisplatin-induced levels of active, phosphorylated ATM (Supplementary Fig. S2A). Combined treatment with ATMi and cisplatin prevented the cisplatin-induced decrease in *BRCA1* expression (Fig. 2D). Consistent with the association between decreased *BRCA1* levels and increased %ALDH+ cells, combining ATMi with cisplatin prevented the cisplatin-induced increase in %ALDH+ cells (Fig. 2E). ATMi itself did not alter the %ALDH+ cells in OVCAR5 cells but decreased the %ALDH+ cells compared to DMSO treated cells in OVSAHO cells (Fig. 2E).

Next, we transiently transfected cells with a plasmid that drives expression of *BRCA1* using an exogenous promoter lacking normal *BRCA1* regulatory regions, including the promoter CpG island (CpGi-null BRCA1; Supplementary Fig. S2B). Transfection of CpGi-null BRCA1 into OVCAR5 and OVSAHO cells resulted in higher *BRCA1* expression than in untreated EV cells even after cisplatin treatment (Fig. 2F). BRCA1 protein levels were also higher in untreated CpGi-null BRCA1 OVCAR5 cells compared to untreated EV cells (Fig. 2G). BRCA1 protein levels decreased in cisplatin-treated CpGi-null BRCA1 cells compared to untreated CpGi-null BRCA1 cells but remained higher than cisplatin treated EV cells (Fig. 2G).

Next, we sought to determine how maintaining BRCA1 levels effects platinum-induced OCSC enrichment. Even though cisplatin increased the %ALDH+ cells in EV as expected, there was no increase in %ALDH+ cells after cisplatin treatment in CpGi-null BRCA1 cells (Fig. 2H). Collectively, these data demonstrate that alteration of BRCA1 levels independent of DNA damage has limited effect on %ALDH+ cells and maintaining *BRCA1* expression prevents the platinum-induced increase in %ALDH+ cells.

### Decitabine treatment abrogates the cisplatin-induced increase in %ALDH+ cells

DNA hypomethylating agents like decitabine (DAC) have been shown to re-sensitize platinum-resistant OC cells to platinum (34). Here, we used low dose DAC to determine the role of DNA methylation in the cisplatin-induced OCSC enrichment. DAC treatment resulted in similar or lower %ALDH+ cells as compared to untreated OVCAR5 and OVSAHO cells, respectively (Fig. 3A). Dual treatment with DAC and cisplatin blocked the cisplatin-induced increase in %ALDH+ cells with the %ALDH+ cells being similar to untreated and/or DAC only treated cells (Fig. 3A). To determine the role of low dose DAC and cisplatin dual treatment on OCSC survival, we examined the ability of pretreated cells to grow as spheroids in stem cell media. DAC pretreated spheroids had similar viability as spheroids generated from non-pretreated cells (Fig. 3B). Dual pretreatment of DAC and cisplatin abrogated the cisplatin-induced spheroid formation and increase in viable cells. Altogether, these data demonstrate that low dose DAC prevents the platinum-induced enrichment of OCSCs.

**Fig 3.**
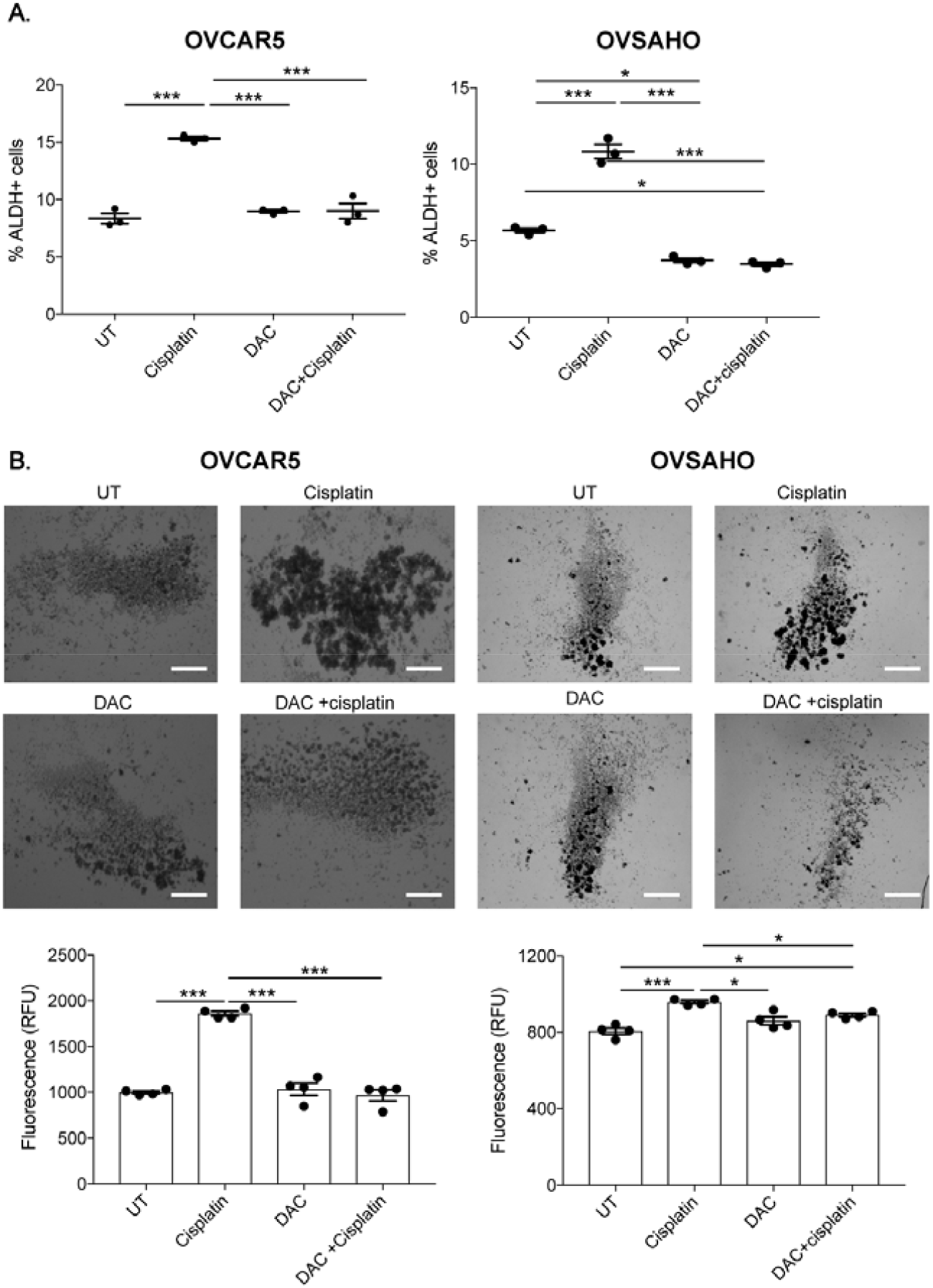
Decitabine treatment abrogates the cisplatin-induced increase in %ALDH+ cells. **A)** Percentage of ALDH+ cells following mock (UT), cisplatin (16h, IC50, OVCAR5: 12 μM, OVASHO: 4 μM), decitabine (DAC; 48h, 100 nM) or DAC + cisplatin treatment. N=3. **E)** Images of spheroids after pretreatment with mock (UT), cisplatin (3h, ½ IC50, OVCAR5: 6 μM, OVSAHO: 2 μM), DAC (48h, 100 nM) or DAC + cisplatin. Scale bar = 500 μm. Graph depicts relative fluorescence units (RFU) of CytoCalcein Violet 450 stain. N=4. For all panels, graphs indicate mean +/- SEM, \**P*<0.05, \*\**P*<0.001, \*\*\**P*<0.0001.

### G2/M cell cycle arrest is associated with an increase in %ALDH+ cells

Because platinum induces cell cycle arrest (35), we studied if cell cycle arrest is related to cisplatin-induced OCSC enrichment. In untreated cells, a higher percentage of ALDH+ cells were in the G2/M phase of the cell cycle than ALDH-cells (Fig. 4A, Supplementary Fig. S3A), consistent with a prior study (35). Additionally, cisplatin treatment resulted in an expected increase in cells in G2/M for both ALDH- and ALDH+ cells (Fig. 4B).

**Fig 4.**
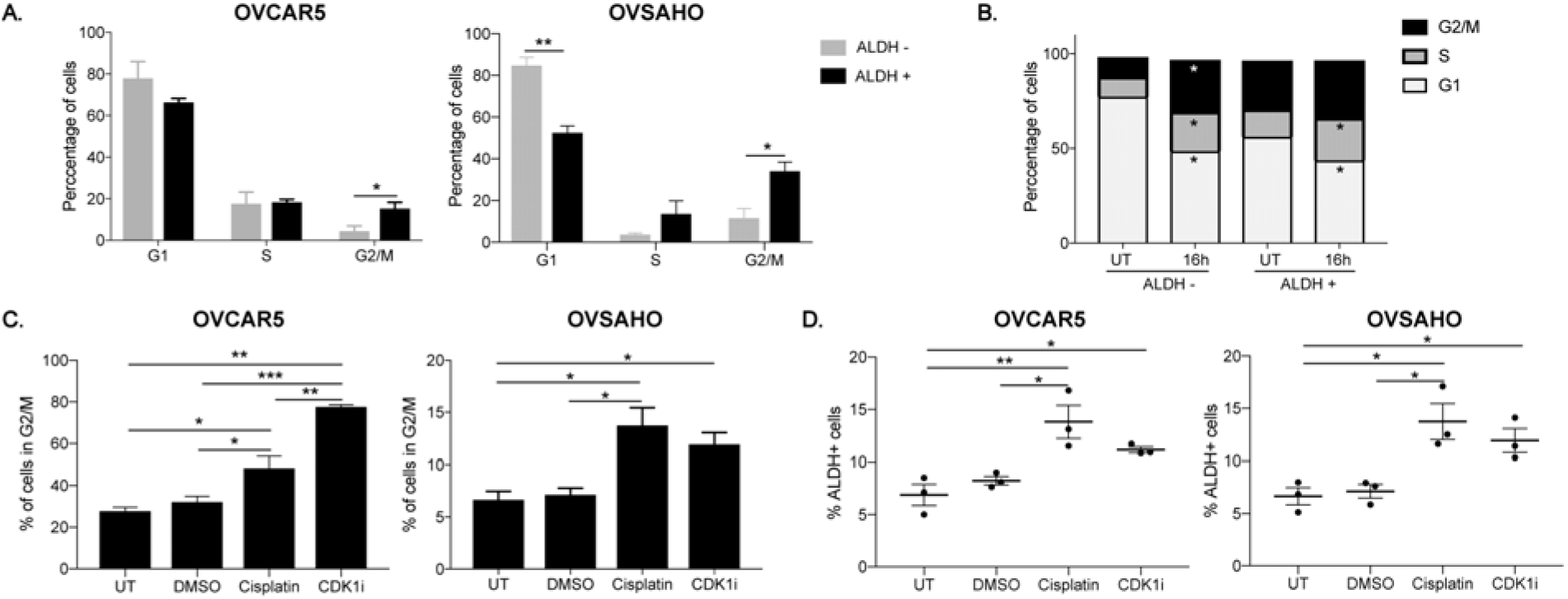
G2/M cell cycle is associated with an increase in %ALDH+ cells. **A)** Percentage of ALDH+ and ALDH-cells in G1, S and G2/M phases of the cell cycle. N=3. **B)** Percentage of ALDH+ and ALDH-OVCAR5 cells in G1, S and G2/M phases of the cell cycle after mock (UT) or cisplatin treatment (16h, IC50, 12 μM). **C)** Percentage of cells in G2/M after mock (UT), DMSO, cisplatin (16h, IC50, OVCAR5: 12 μM, OVSAHO: 4 μM) or CDK1 inhibitor (RO-3306, 16h, 9 μM) treatment. N=3. **D)** Percentage of ALDH+ cells treated as in C. N=3. For all panels, graphs indicate mean +/- SEM, \**P*<0.05, \*\**P*<0.001, \*\*\**P*<0.0001.

To determine if G2/M arrest is important for the cisplatin-induced increase in %ALDH+ cells, we induced G2/M arrest independent of platinum treatment through cyclin-dependent kinase 1 inhibition (CDK1i). CDK1i increased the percentage of cells in G2/M compared to controls to a level that was similar to or higher than levels after cisplatin treatment in OVSAHO and OVCAR5 cells, respectively (Fig. 4C, Supplementary Fig. S3B). Comparably to cisplatin, CDK1i resulted in a significant increase in %ALDH+ cells compared to untreated (Fig. 4D). Additionally, unlike cisplatin treatment, DNA damage was not increased by CDK1i (measured by γH2AX levels, Supplementary Fig S3C), suggesting that G2/M arrest alone can increase %ALDH+ cells.

### NAMPT inhibition abrogates cisplatin-induced enrichment of ALDH+ cells

A key co-factor for ALDH activity is NAD^+^ (36), and increased levels of NAD^+^ and NAMPT have been shown to promote cancer cell survival and reported to drive platinum-induced increase in senescence-associated OCSCs (13, 17, 37). We observed that NAD^+^ levels increased after 16h cisplatin treatment (Fig. 5A). Additionally, *NAMPT* expression increased after 16h cisplatin treatment in OVCAR5 (platinum-sensitive and - resistant), OVSAHO, OVCAR3, PEO1 and COV362 (Fig. 5B, Supplementary Fig. S3D). To determine if blocking the cisplatin-induced increase in NAD^+^ prevented the increase in %ALDH+ cells, we treated cells with a NAMPT inhibitor (NAMPTi) (38). Dual treatment with NAMPTi and cisplatin prevented the cisplatin-induced increase in NAD^+^ levels with NAD^+^ levels being similar to controls (Fig. 5C). NAMPTi treatment alone in OVCAR5 and OVSAHO cells decreased or maintained %ALDH+ cells relative to controls, respectively (Fig. 5D). Combined treatment of NAMPTi and cisplatin prevented the cisplatin-induced increase in %ALDH+ cells, with the %ALDH+ cells in the dual treated samples being decreased or similar to controls in OVCAR5 and OVSAHO, respectively. Further, dual pretreatment with NAMPTi and cisplatin abrogated cisplatin-induced spheroid formation and increased the number of viable cells (Fig. 5E). Dual pretreated cells had similar or lower viability as spheroids generated from DMSO or NAMPTi pretreated cells, respectively. These data demonstrated that cisplatin induced an increase in NAD^+^ levels through increased *NAMPT* expression and NAMPTi abrogated the cisplatin-induced OCSC enrichment.

**Fig 5.**
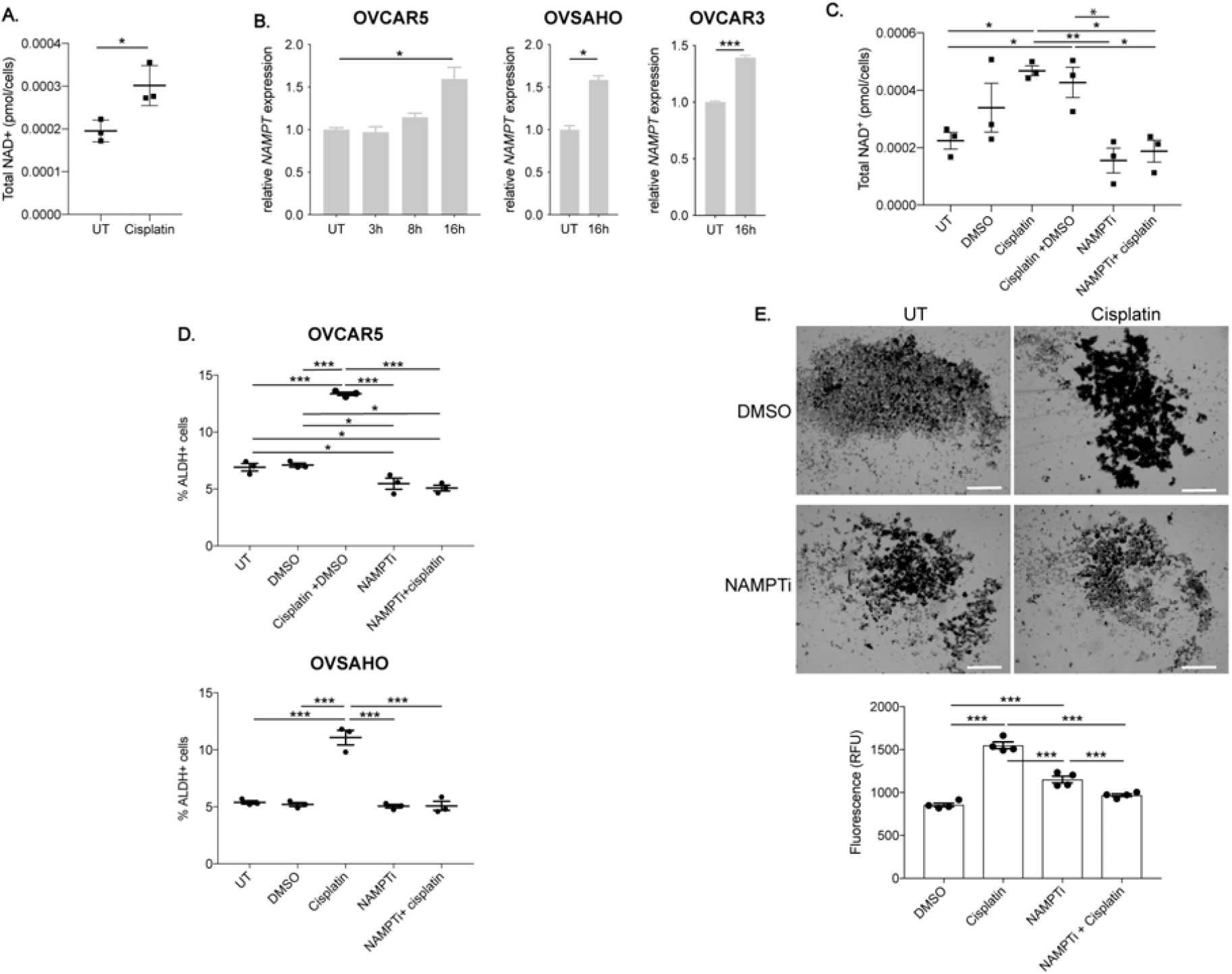
NAMPT inhibition abrogates cisplatin-induced enrichment of ALDH+ cells. **A)** Total NAD^+^ (pmol/cells) in OVCAR5 cells after mock (UT) or cisplatin (16h, IC50, 12 μM) treatment. N=3. **B)** Relative *NAMPT* RNA expression after mock or cisplatin (IC50 dose, OVCAR5: 12 μM, OVSAHO: 4 μM, OVCAR3: 15 μM) treatment. N=3. **C)** Total NAD^+^ (pmol/cells) in OVCAR5 cells after mock (UT), cisplatin (16h, IC50, 12 μM), NAMPTi (STF-118804, 6h, 50 nM), cisplatin + DMSO or cisplatin + NAMPTi treatment. N=3. **D)** Percentage of ALDH+ cells after treatment as in C. N=3. **E)** Images of spheroids after pretreatment with mock (DMSO), cisplatin (3h, ½ IC50, 6 μM), NAMPTi (6h, 50 nM), or NAMPTi + cisplatin treatment. Scale bar = 500 μm. Graph depicts relative fluorescence units (RFU) of CytoCalcein Violet 450 stain. N=4. For all panels, graphs indicate mean +/- SEM, \**P*<0.05, \*\**P*<0.001, \*\*\**P*<0.0001.

### Cisplatin treatment induces two separate pathways to increase %ALDH+ cells

To further explore how decreased *BRCA1* expression, increased NAD^+^ levels and G2/M arrest are interconnected during cisplatin-induced OCSC enrichment, we assayed *BRCA1* and *NAMPT* expression after CDK1i, BRCA1 overexpression and DAC treatment. Following CDK1i (shown to increase the %ALDH+ cells, Fig. 4D), *BRCA1* and *NAMPT* expression increased relative to controls (Supplementary Fig. S4A,B), suggesting that G2/M cell cycle arrest and changes in *NAMPT* but not *BRCA1* expression are connected. Furthermore, *NAMPT* expression was elevated by transfection of either EV or CpGi-null BRCA1 compared to non-transfected, untreated cells and *NAMPT* expression in transfected cells was not further increased by cisplatin (Supplementary Fig. S4C). Cisplatin treatment increased *NAMPT* expression as expected in non-transfected cells; however, in OVSAHO cells, cisplatin treatment increased *NAMPT* expression in both EV and CpGi-null BRCA1 compared to the respective untreated controls (Supplementary Fig. S4C). NAD^+^ levels increased in cisplatin treated EV and CpGi-null BRCA1 cells compared to untreated controls, although to a lesser extent than in non-transfected cells (Supplementary Fig. S4D). Similarly, DAC treatment alone or in combination with cisplatin elevated *NAMPT* expression compared to untreated, and the level was similar to (or higher) than cells treated with cisplatin alone (Supplementary Fig. S4E). These data suggested that even though maintaining BRCA1 expression blocks the cisplatin-induced increase in %ALDH+ cells (Fig. 2D, 3D), cisplatin treatment still increases *NAMPT* and NAD^+^ levels.

As BRCA1 overexpression has been previously connected to an increase of cells in the G2/M phase of the cell cycle (39), we determined the effect of CpGi-null BRCA1 expression on the cell cycle. Transfection alone increased cells in G2/M, with CpGi-null BRCA1 cells having higher levels than EV cells, which were higher than non-transfected cells (Supplementary Fig. S4F). Cisplatin treatment increased cells in G2/M for all sample types relative to their respective untreated controls, again with CpGi-null BRCA1 cells having the highest percentage of cells in G2/M (Supplementary Fig. S4F). Altogether, these data indicated that the platinum-induced G2/M arrest and the associated change in *NAMPT* expression and NAD+ levels still occurred even with sustained BRCA1 expression and loss of the platinum-induced increase in %ALDH+ cells.

### Dual DNMT and NAMPT inhibitor treatment abrogates the cisplatin-induced increase in %ALDH+ cells

The above observations indicated that DNA methylation and increased NAD+ levels contribute to the cisplatin-induced enrichment of OCSCs. Although inhibiting either alteration alone with DAC (Fig. 3) or NAMPTi (Fig. 5) abrogated the cisplatin-induced OCSC enrichment, we hypothesized that combining very low dose treatment of the two inhibitors would impact both aspects of the platinum-induced changes and further block the cisplatin-induced increase in %ALDH+ cells.

First, DAC and NAMPTi were used alone to determine concentrations that had minimal to no effect on the cisplatin-induced increase in %ALDH+ cells (Supplementary Fig. S5A, B). Then, we sought to determine if combining selected lower doses of each drug would prevent the cisplatin-induced increase in %ADLH+ cells. Combination treatment of very low dose DAC with low dose NAMPTi and cisplatin prevented the cisplatin-induced increase in %ALDH+ cells and resulted in similar %ALDH+ cells as DMSO treated cells, while the individual treatments had no effect (Fig. 6A). Furthermore, the combined treatment was calculated to be synergistic (CI< 1; (40)) in ability to block the cisplatin-induced increase in %ALDH+ cells (Fig. 6B).

**Fig 6.**
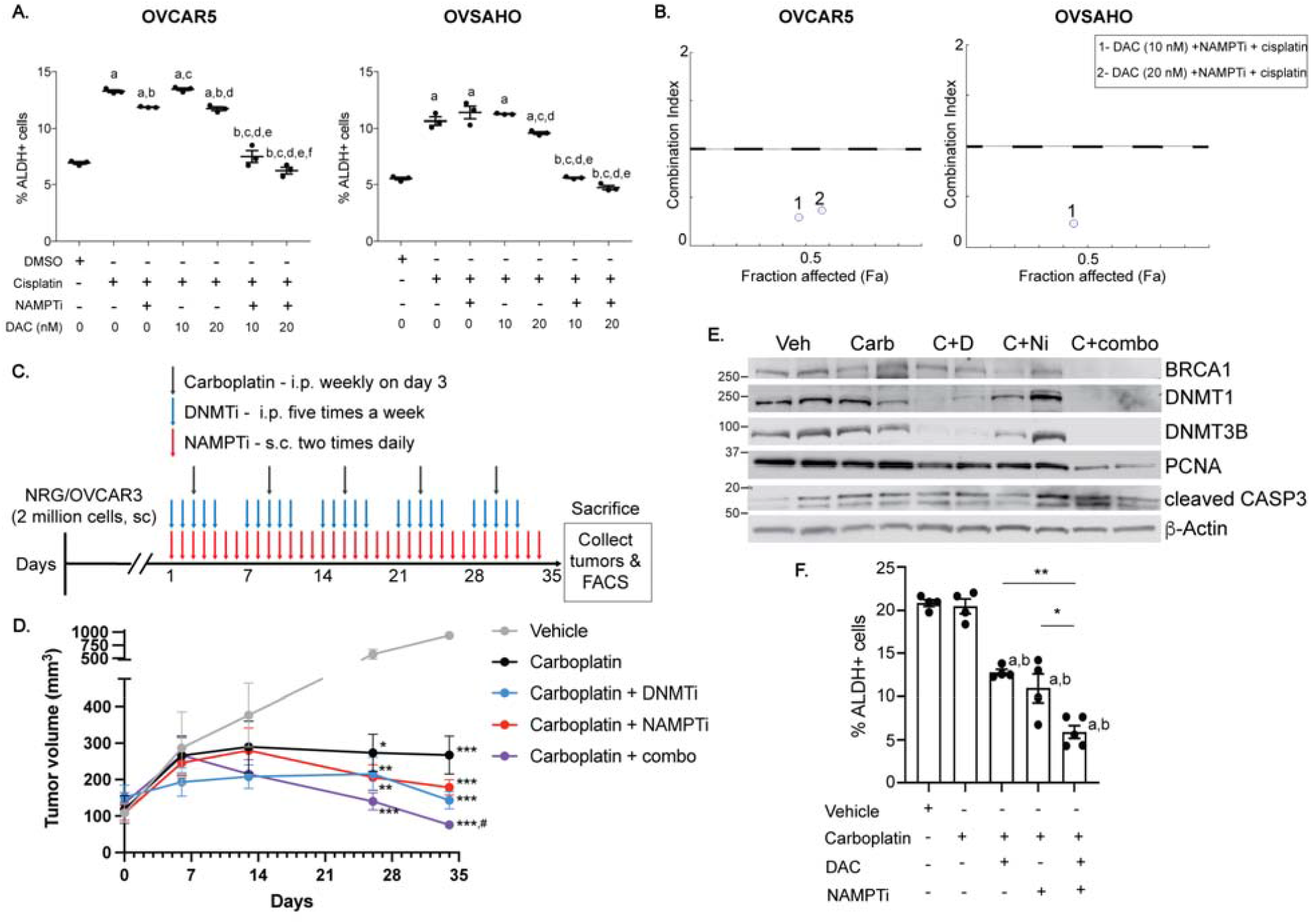
Dual DNMTi and NAMPTi treatment abrogates cisplatin-induced OCSC enrichment. **A)** Percentage of ALDH+ cells after treatment with mock (DMSO), cisplatin (16h, IC50, OVCAR5: 12 μM, OVSAHO: 4 μM), DAC (decitabine; 48h, 10 nM or 20 nM) + cisplatin, NAMPTi (6h, 12.5 nM) + cisplatin or NAMPTi +DAC + cisplatin. N=3. *a* indicates *P*<0.05 for comparison to DMSO treated. *b* indicates comparison to cisplatin treated. *c* indicates comparison to NAMPTi+ cisplatin treated. *d* indicates comparison to DAC (10 nM) + cisplatin. *e* indicates comparison to DAC (20 nM) + cisplatin. *f* indicates comparison to NAMPTi + DAC (10 nM) + cisplatin. **B)** Combination index plot for cells treated as described in A; x-axis represents fraction affected (Fa) and y-axis represents combination index. Combinates beneath the dashed line are synergistic. **C)** Treatment scheme for xenograft experiment (carboplatin (25 mg/kg weeks 1 and 2; 50 mg/kg weeks 3,4, and 5), DAC (0.1 mg/kg), NAMPTi (6.25 mg/kg)). **D)** Tumor volume determined by digital caliper measurement over 5 weeks of treatment. N=4-5 mice per group. \**P* relative to vehicle, #*P* relative to carboplatin only. **E)** Western blot of xenograft protein lysates. **F)** Percentage of ALDH+ cells in single cells dissociated from xenografts. *a* indicates *P*<0.001 for comparison to vehicle. *b* indicates *P*<0.001 for comparison to carboplatin only. For all panels, graphs depict mean +/- SEM. \**P*<0.05, \*\**P*<0.001, \*\*\**P*<0.0001.

To confirm the effect of very low dose DAC and low dose NAMPTi on cisplatin-induced enrichment of OCSCs, the ability of pretreated cells to grow as spheroids in stem cell media was examined. Cells pretreated with individual very low doses of DAC or low dose NAMPTi and cisplatin were similar to cisplatin only pretreated cells (Supplementary Fig. S5C). Importantly, combination pretreatment of very low dose DAC, low dose NAMPTi and cisplatin prevented cisplatin-induced spheroid formation and increased viability.

To determine if DAC and NAMPTi alter tumorigenesis and OCSC enrichment *in vivo*, we established OVCAR3 flank xenografts in immunodeficient mice. Randomized mice received vehicle, carboplatin alone or carboplatin with low dose DAC, low dose NAMPTi or low dose DAC+ low dose NAMPTi (Fig. 6C for dosing schedule). As expected, carboplatin alone or in combination with the inhibitors reduced tumor volume starting at week 4 of treatment (Fig. 6D). After 5 weeks of treatment, tumors from the carboplatin+DAC+NAMPTi group were significantly smaller than those from the carboplatin alone group. Conversely, tumor size in DAC or NAMPTi with carboplatin treatment was not different than carboplatin alone. In all groups of mice treated with carboplatin, body weight by the end of the study was reduced but not statistically different compared to the vehicle only, and no additional effect on body weight was seen in the combined DAC and NAMPTi group (violet line; Supplementary Fig. S5D), suggesting that the low doses of these inhibitors used were well tolerated. As expected, DNMT1 and DNMT3B levels were decreased in tumors from DAC-treated mice (carboplatin+DAC and carboplatin+combo) compared to vehicle or carboplatin only treated mice (Fig. 6E). Levels of the proliferation marker PCNA were decreased in tumors from carboplatin+DAC-and carboplatin+NAMPTi-treated mice compared to vehicle or carboplatin only treated mice (Fig. 6E). Furthermore, the lowest levels of PCNA were seen in tumors from carboplatin+DAC+NAMPTi-treated mice, indicating that these tumors had the lowest level of proliferative cells. There was no obvious difference in cleaved caspase 3 or BRCA1 levels between the treatment groups, which is likely because the samples were collected 6 days after the last carboplatin treatment.

To determine the effect of the inhibitors on OCSC enrichment, we performed ALDEFLUOR assays on dissociated tumor cells. In contrast to previous studies (2, 23, 41), carboplatin alone did not enrich for ALDH+ cells in the xenografts when compared to vehicle (Fig. 6F). Nevertheless, treatment with carboplatin and DAC or NAMPTi alone or carboplatin+DAC+NAMPTi reduced the %ALDH+ cells in the xenografts as compared to vehicle or carboplatin treatment alone. Importantly, the %ALDH+ cells in tumors from carboplatin+DAC+NAMPTi treated mice was lower than in tumors from mice treated with carboplatin and either inhibitor alone (Fig. 6F).

## Discussion

Our study identifies two alterations that drive chemotherapy-induced OCSC enrichment, one marked by reduced *BRCA1* expression and the other by increased NAD^+^ levels. BRCA1 plays a role in ICL repair as well as other parts of the DDR (11). ICL accumulation following BRCA1 depletion results in dedifferentiation of mammary epithelial cells to a more primitive, mesenchymal state (29). Our ATMi and BRCA1 KD results demonstrate that the DDR after platinum treatment is important for the proposed mechanism. We speculate that the platinum-induced decrease in BRCA1 levels leads to persistent ICLs or alternative DNA repair pathway activation, which is required for but not sufficient for OCSC enrichment. Expression of CpGi-null BRCA1 maintained *BRCA1* expression levels at or above the levels of untreated cells even after cisplatin treatment and blocked the platinum-induced OCSC enrichment. This finding suggests a “threshold effect” for BRCA1 levels: BRCA1 levels at or above baseline prevent enrichment of ALDH+ cells, and OCSC enrichment occurs when BRCA1 levels fall below baseline in the presence of an activated DDR.

Interestingly, even though cisplatin resistant cells had a higher baseline percentage of ALDH+ cells than parental sensitive cells, *BRCA1* expression levels were higher in the resistant cells. Furthermore, platinum treatment *in vivo* did not alter BRCA1 protein levels in tumors collected at the end-of-study. Our findings suggest that platinum-induced *BRCA1* repression is transient but required for the platinum-induced increase in %ALDH+ cells. We hypothesize that the platinum-induced decrease in BRCA1 levels serves to initiate the increase in %ALDH+ cells and once ALDH+ cells are enriched, but low levels of BRCA1 levels are not necessary to maintain the %ALDH+ increase. This hypothesis is consistent with findings that decreasing *BRCA1* expression increased stemness of mammary epithelial cells and that the increase in stemness was maintained even after *BRCA1* expression was restored (29).

In addition to the DDR-dependent changes in *BRCA1* expression, we demonstrate that platinum induces a concurrent increase in NAD^+^, and altering this metabolic pathway is also required for platinum-induced OCSC enrichment. Because NAD^+^ is a cofactor for ALDH, the platinum-induced increase in NAD^+^ likely drives the increased ALDH activity of OCSCs. Cellular NAD^+^ production has been linked to CDK1i and G1 and G2 cell cycle phases (42, 43), and we show that CDK1i induces OCSC enrichment that is associated with an increase in *NAMPT* expression (Supplementary Fig. S5B). Consistent with BRCA1’s role in G2/M arrest (44), CpGi-null BRCA1 transfection increases the percentage of cells in G2/M regardless of cisplatin treatment (Supplementary Fig. S5E) but blocks platinum-induced OCSC enrichment. Together these findings suggest that G2/M arrest leads to increased *NAMPT* expression and NAD^+^ levels that contribute to OCSC enrichment; however, for platinum-induced OCSC enrichment to occur, these changes must occur in conjunction with decreased levels of BRCA1. While platinum treatment did not alter ALDH1A expression in our system (Supplementary Figure S1C), others have found that cisplatin treatment increases ALDH expression (45, 46). There are 19 human ALDH isoforms and several signaling pathways regulate ALDH expression (47). Cisplatin may have different effects on ALDH expression based on the time point examined after cisplatin treatment, the major isoform of ALDH expressed, and signaling pathways activated in the model system being used.

Loss of BRCA1 induces metabolic reprogramming through the nicotinamide N-methyltransferase (NNMT) pathway in OC cells (48). Furthermore, loss of *BRCA1* expression by mutation or methylation is correlated with increased NAMPT dependent NAD+ production in OC patient tissue and cell lines (16). Therefore, we hypothesized that decreased BRCA1 and increased NAD^+^ levels may be mechanistically connected. However, even though sustained expression of *BRCA1* in CpGi-null BRCA1 transfected cells blocked the platinum-induced increase in %ALDH+ cells, NAD^+^ levels still increased (Supplementary Fig. S4D). Importantly, in *BRCA1* mutant COV362 cells, cisplatin increased *NAMPT* expression but not enrichment of ALDH+ cells (Supplementary Fig. S3D,S1B). We suggest that the platinum-induced decrease in BRCA1 and increase in *NAMPT* and NAD^+^ levels occur as distinct parallel pathways which independently contribute to platinum-induced OCSC enrichment. In the present study, we incorporated several different inhibitors, including CDK1, ATM, NAMPT and DNMTs inhibitors. While highly specific, these inhibitors may also impact broad cellular processes such as metabolism, cell cycle progression, and apoptosis that could confound the interpretation of our results. Our future studies will further investigate how platinum-induced DDR and metabolic alterations result in OCSC enrichment and expand the therapeutic relevance of our findings by including patient derived and orthotopic models.

Clinically, DNMTi have been used to re-sensitize chemotherapy resistant OC cells to chemotherapy (34, 49-52). Our study extends these findings by demonstrating that DNMTi may also be beneficial when used in combination with neoadjuvant/adjuvant chemotherapy in OC patients where they have the potential to block platinum-induced OCSC enrichment and establishment of platinum resistance. NAMPTi have also been clinically tested in advanced hematological and solid malignancies (53, 54). However, dose limiting toxicities were a significant problem and objective tumor remission was not observed. The finding that low dose NAMPTi combined with very low dose DNMT inhibitors and platinum was both an effective treatment scheme to reduce OCSC enrichment and well tolerated in vivo is an encouraging new finding. Even though we and others have previously observed platinum-induced enrichment of OCSCs *in vivo* (23, 41), that finding was not replicated in our study (Fig. 6F). We believe that this may have been a technical issue due to the mouse strain used or the study length, which was 3 weeks in the previous studies compared to 5 weeks in the present study. However, this result could also suggest that DAC and NAMPTi are working through a different mechanism *in vivo*. The *in vivo* doses of DAC and NAMPTi were based on previously published studies (38, 55, 56). Because both inhibitors still influenced OCSC enrichment when combined separately with carboplatin, it is possible that doses of these inhibitors could be reduced even further when combined *in vivo* to minimize toxicity while still maintaining efficacy. Altogether our findings suggest the combining epigenetic and metabolic inhibitors will prevent onset of platinum resistance in wildtype *BRCA1*, platinum-sensitive HGSOC. Using lower concentrations of these inhibitors in combination may reduce off-target cytotoxic effects and make the treatment more tolerable to patients.

## Acknowledgments

We thank the Indiana University Flow Cytometry Core Facility for their assistance. This research was funded in part by the Ovarian Cancer Research Alliance (grant number 458788 to HMOH and KPN), the Ovarian Cancer Alliance of Greater Cincinnati (to KPN) and through the IU Simon Comprehensive Cancer Center P30 Support Grant (P30CA082709-20). SS was supported by the Doane and Eunice Dahl Wright Fellowship generously provided by Ms. Imogen Dahl.

