## Supplemental materials and methods for "Combined epigenetic and metabolic inhibition blocks platinum-induced ovarian cancer stem cell enrichment"

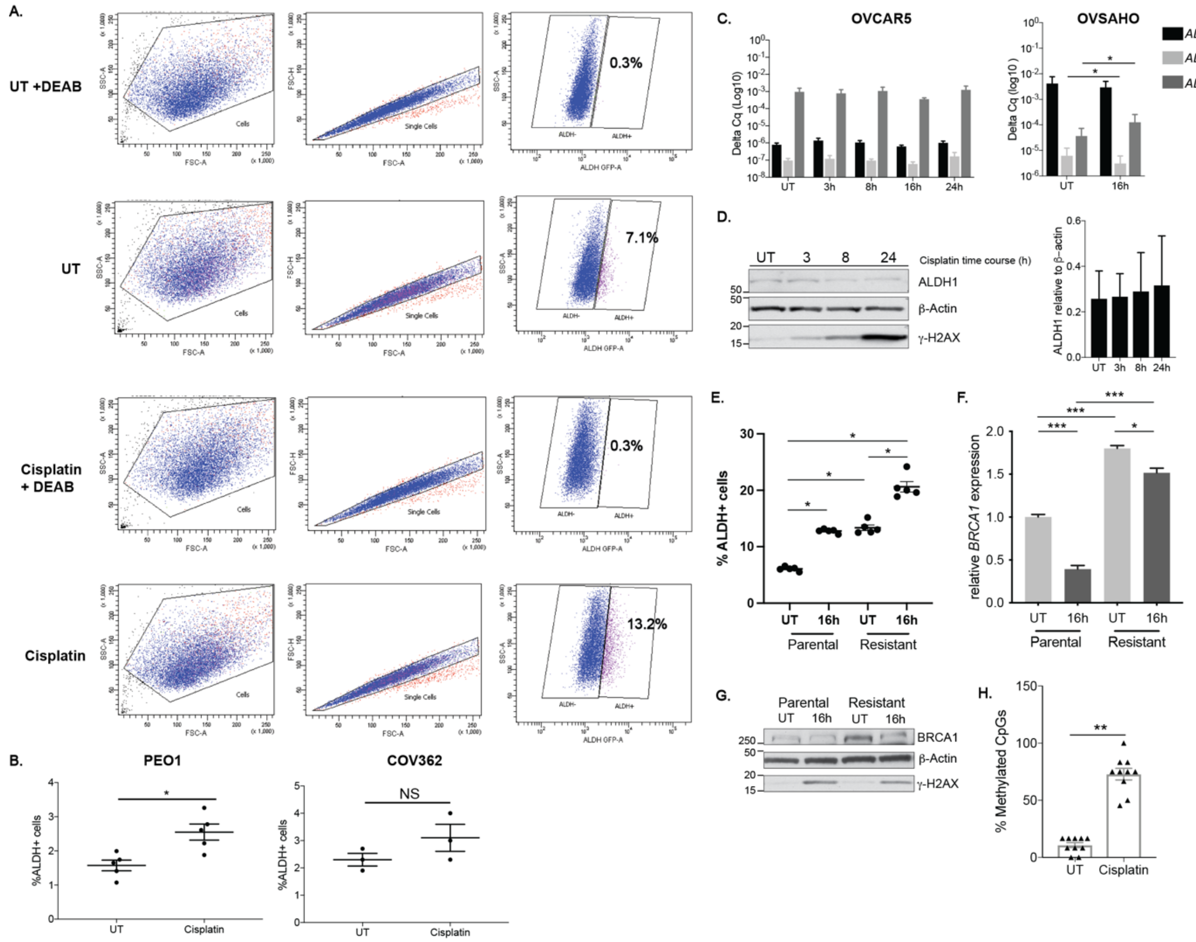


**Supplementary Fig S1: A)** Plots for ALDH+/- percentage in OVCAR5 cells with DEAB treatment (negative control) or ALDEFLUOR assay reagent (ALDH) after mock (UT) or cisplatin treatment (16 h, IC50, 12 ⌈M for OVCAR5 cells). **B)** %ALDH+ cells determined using the ALDEFLUOR assay after mock (UT) or cisplatin (16 h, IC50, PEO1: 12.84 ⌈M, COV362: 13.57 ⌈M) for indicated cell lines. **C)** *ALDH1A* isoform RNA expression in indicated cell lines after mock or cisplatin treatment (IC50, OVCAR5: 12 ⌈M, OVSAHO: 4 ⌈M) for the indicated time points. Graphs depict mean delta Cq of isoform relative to *PPIA*. **D)** Western blot and relative densitometry of OVCAR5 whole cell lysates after treatment as in C for the indicated time points. **E)** %ALDH+ cells determined using the ALDEFLUOR assay after mock (UT) or cisplatin (16 h, 12 mM) for parental and resistant OVCAR5 cells.

**F)** Relative BRCA1 RNA expression following treatment as in E. **G)** Western blot of OVCAR5 parental and resistant whole cell lysates following treatment as in E. **H)** Percentage of methylated CpGs per clone using data presented in Figure 1F. Graphs depict mean + SEM. N=10. For all panels, N=3 except where indicated otherwise, graphs indicate mean +/- SEM, **P*<0.05, ***P*<0.001, ****P*<0.0001,

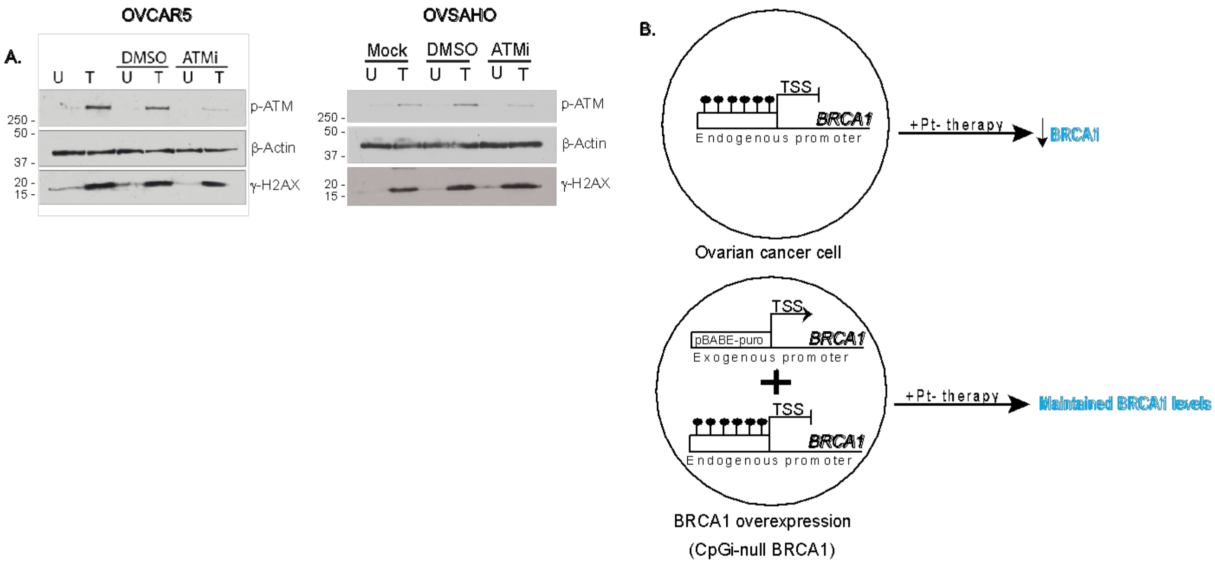


**Supplementary Fig S2: A)** Western blot of whole cell lysates after mock (UT), DMSO or ATMi (16 h, 15 ⌠M) without (U) or with (T) cisplatin treatment (16 h, IC50, OVCAR5: 12 ⌠M, OVSAHO: 4 ⌠M) in indicated cell lines. **B)** Schematic diagram for CpGi-null BRCA1 transfection.

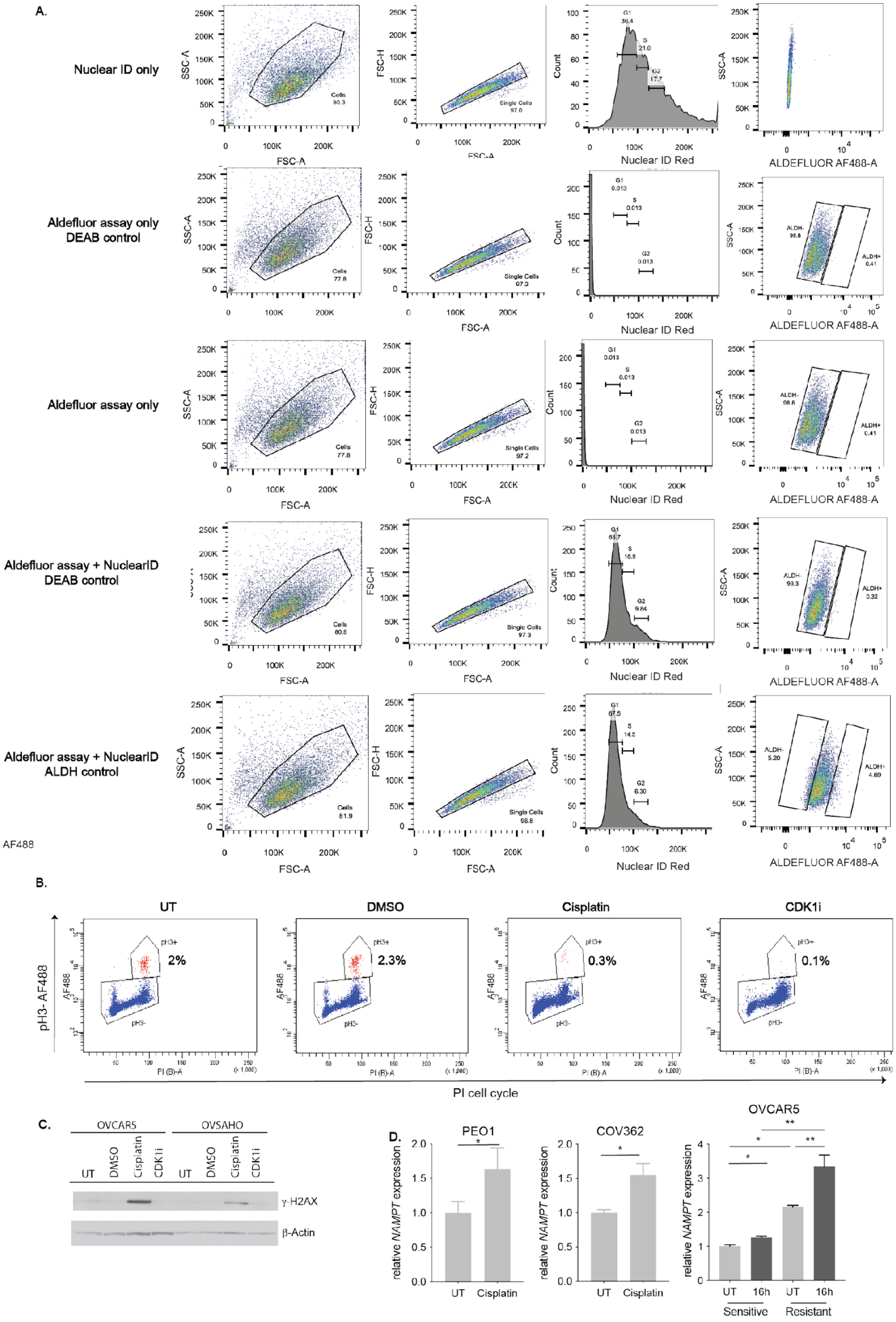
A.
Nuclear ID only
Aldefluor assay only DEAB control
Aldefluor assay only
Aldefluor assay + NuclearID DEAB control
Aldefluor assay + NuclearID ALDH control
AF488
SSC-A
FSC-A
FSC-H
Count
Nuclear ID Red
ALDEFLUOR AF488-A
Cells
Single Cells
G1
S
G2
ALDH-
ALDH+
B.
UT
DMSO
Cisplatin
CDK1i
pH3- AF488
pH3+
pH3-
2%
2.3%
0.3%
0.1%
PI cell cycle
C.
OVCAR5
OVSAHO
UT
DMSO
Cisplatin
CDK1i
γ-H2AX
β-Actin
D.
PEO1
COV362
OVCAR5
relative NAMPT expression
UT
Cisplatin
16h
Sensitive
Resistant

**Supplementary Fig S3: A)** Plots for ALDH+/- percentage with DEAB treatment (negative control) or ALDEFLUOR assay reagent (ALDH) with Nuclear ID Red stain in OVCAR5 cells. **B)** Plots for phosphorylated serine 10 histone H3 +/- cells percentage after mock (UT), DMSO, cisplatin (16 h, IC50, 12 ⌠M) or CDK1 inhibitor (16 h, 9 μM) treatment in OVCAR5 cells. X: axis- PI stain, Y axis: pH3 with secondary conjugated with AF-488. **C)** Western blot of whole cell lysates after mock (UT), DMSO, cisplatin (16 h, IC50, OVCAR5: 12 μM, OVSAHO: 4 μM) or CDK1 inhibitor (16 h, 9 μM) treatment in indicated cell lines. **D)** *NAMPT* RNA expression after mock or cisplatin treatment (16 h, IC50, PEO1: 12.84 μM, COV362: 13.57 μM, OVCAR5: 12 μM) for the indicated time points and cell lines. Graphs depict mean + SEM. N=3, *$P<0.05$, **$P<0.001$, ***$P<0.0001$.

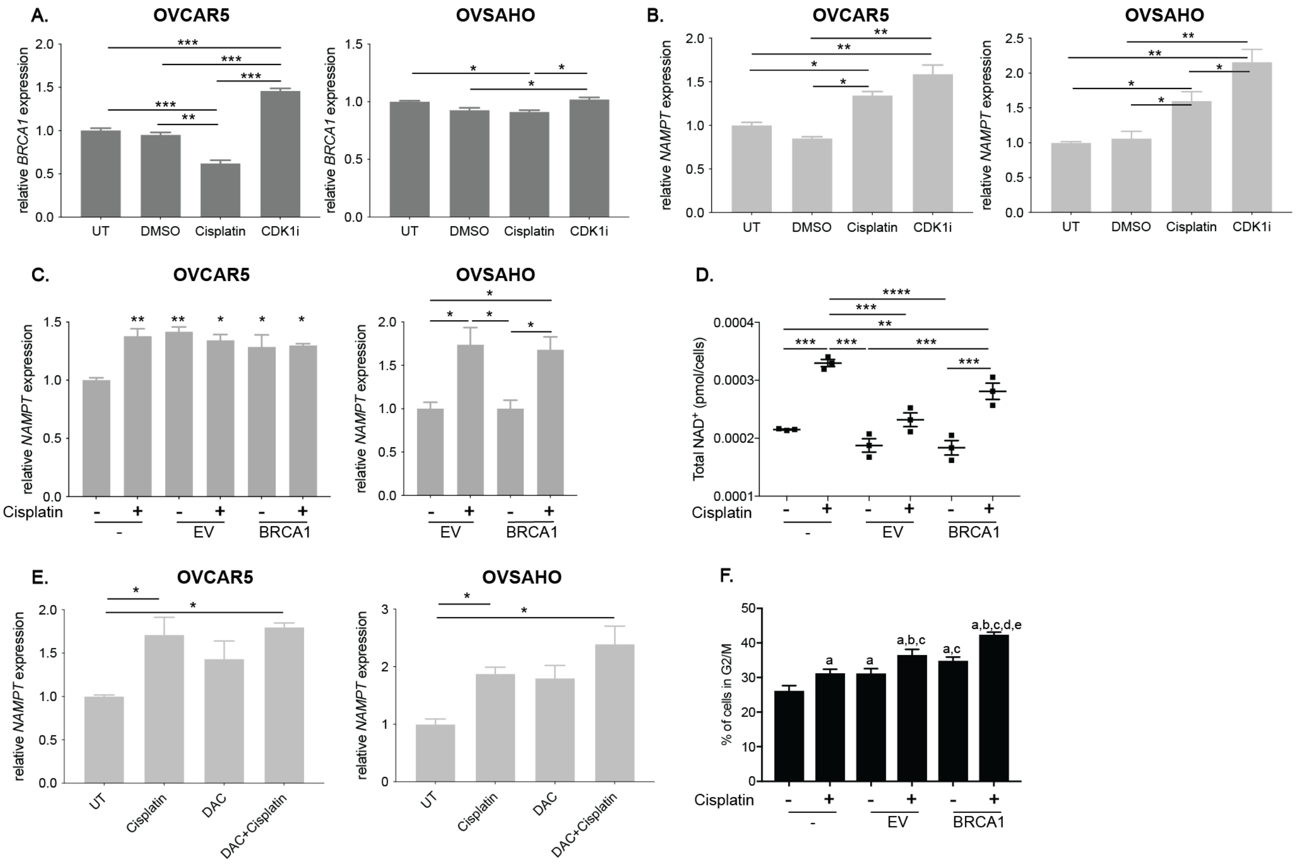


**Supplementary Fig. S4. A)** Relative *BRCA1* RNA expression after treatment with mock (UT), DMSO, cisplatin (16 h, IC50, OVCAR5: 12 ⌈M, OVSAHO: 4 ⌈M) or CDK1 inhibitor (16 h, 9 ⌈M) treatment in indicated cell lines. N=3. **B)** Relative *NAMPT* RNA expression after treatment as in A in indicated cell lines. N=3. **C)** Relative *NAMPT* RNA expression in untransfected (UT) or transfected with mock empty vector (EV) or CpGi-null BRCA1 plasmid and treated with or without cisplatin (16 h, IC50, OVCAR5: 12 ⌈M, OVSAHO: 4 ⌈M) in the indicated cell lines. For OVCAR5, * indicates comparison between the indicated sample type and untransfected control with no treatment. N=3. **D)** Total NAD$^+$ (pmol/cells) in OVCAR5 cells untransfected (UT) or transfected with mock empty vector (EV) or CpGi-null BRCA1 (BRCA1) plasmid and treated with or without cisplatin (16 h, IC50, 12 ⌈M).

N=3. **E)** Relative *NAMPT* RNA expression after mock (UT), cisplatin (16 h, IC50, OVCAR5: 12 ⌈M, OVSAHO: 4 ⌈M), DAC (48 h, 100 nM) or DAC (48 h, 100 nM) + cisplatin (16 h, IC50) treatment in indicated cell lines. N=3. **F)** Percentage of cells in G2/M in OVCAR5 cells untransfected (UT) or transfected with mock empty vector (EV) or CpGi null BRCA1 plasmid and treated as in D. N=3. a indicates comparison between the indicated sample type and untransfected UT sample. b indicates comparison between the indicated sample type and untransfected cisplatin treated sample. c indicates comparison between the indicated sample type and sample transfected with mock (EV). d indicates comparison between the indicated sample type and mock (EV) treated with cisplatin. e indicates comparison between the indicated sample type and sample or sample transfected with CpGi-null promoter BRCA1 (BRCA1) plasmid. For all panels, graphs indicate mean +/- SEM, *$P$<0.05, **$P$<0.001, ***$P$<0.0001, ****$P$<0.00001.

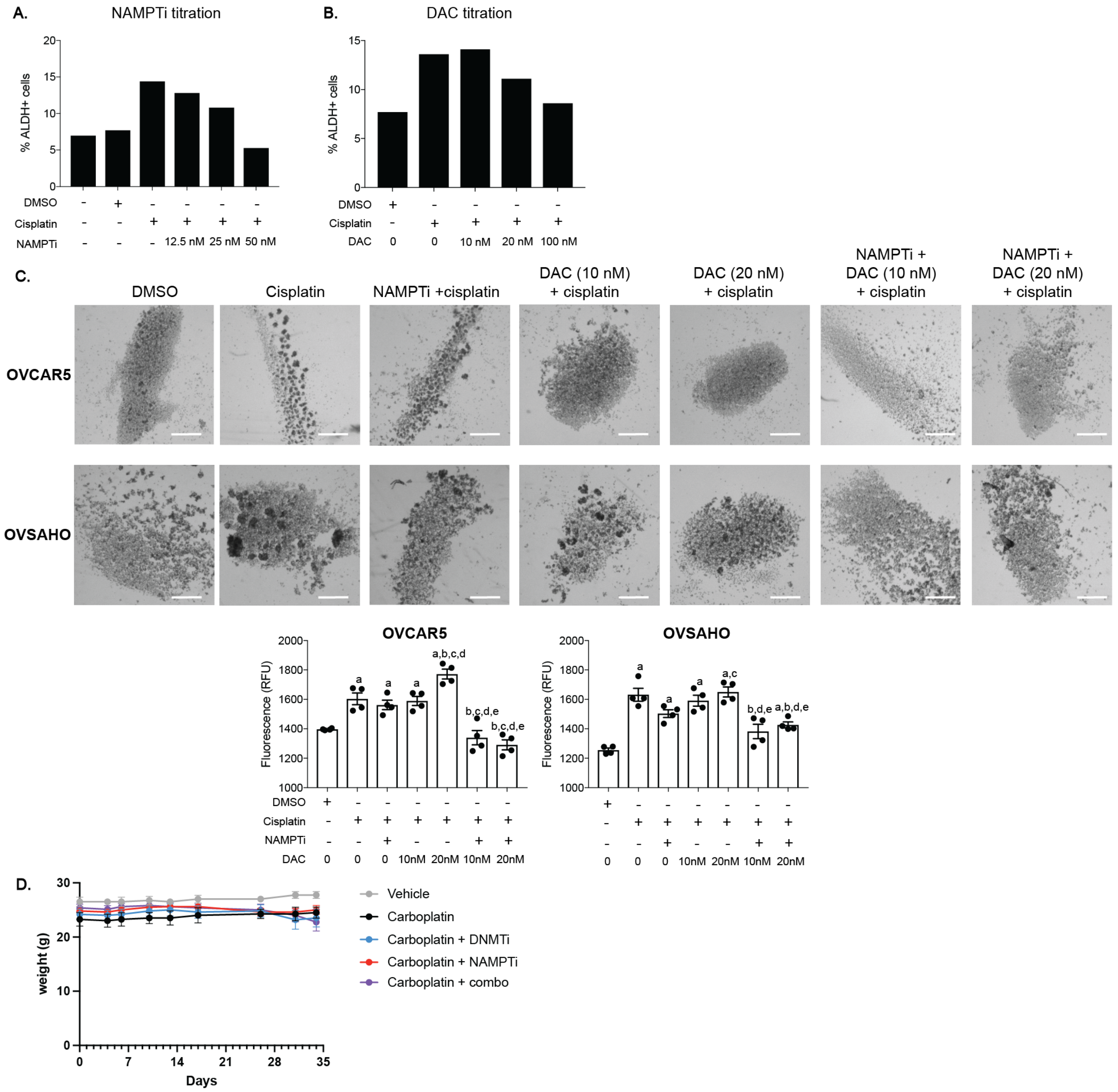


**Supplementary Fig. S5. A)** Percentage of ALDH+ cells using the ALDEFLUOR assay after treatment of OVCAR5 cells with mock (UT or DMSO), cisplatin (16 h, IC50, 12 ⌈M) or NAMPTi (6 h, 12.5 nM, 25 nM or 50 nM) + cisplatin. **B)** Percentage of ALDH+ OVCAR5 cells using the ALDEFLUOR assay after treatment with mock (UT or DMSO), cisplatin (16 h, IC50, 12 ⌈M) or DAC (48 h, 10 nM, 20 nM or 100 nM) + cisplatin. **C)** Images of spheroids after mock (DMSO), cisplatin (3 h, ½ IC50, OVCAR5: 6 ⌈M,

OVSAHO: 2 ⌠M), DAC (48 h, 10 nM or 20 nM) + cisplatin, NAMPTi (6 h, 12.5 nM) + cisplatin treatment or NAMPTi +DAC + cisplatin treated in indicated cell lines. Scale bar = 500 µm. Graph depicts fluorescence intensity (RFU) of CytoCalcein Violet 450 stain from spheroids in indicated cell lines. N=4. Graphs depict mean +/- SEM. a indicates *P*<0.05 for comparison between the indicated sample type and DMSO treated sample. b indicates comparison between the indicated sample type and cisplatin treated sample. c indicates comparison between the indicated sample type and NAMPTi+ cisplatin treated sample. d indicates comparison between the indicated sample type and DAC (10 nM) + cisplatin treated sample. e indicates comparison between the indicated sample type and DAC (20 nM) + cisplatin treated sample. f indicates comparison between the indicated sample type and NAMPTi + DAC (10 nM) + cisplatin treated sample. **D)** Weights of mice in each group over the 35 days of treatment. N=4-5 mice per treatment group. Graphs depict mean +/- SEM.

**Supplementary Table S1. Primers, Taqman assays, and antibodies used.**

| Gene | Forward Sequence (5'-> 3') | Reverse Sequence (5'-> 3') |
|---|---|---|
| *ALDH1A1* | TCCCGTTGGTTATGCTCATTTG | GGAGTTTGCTCTGCTGGTTTGAC |
| *ALDH1A2* | TTGGTTCAGTGTGGAGAAGG | AAAGCTTGCAGGAATGGTTTG |
| *ALDH1A3* | CTTCTGCCTTAGAGTCTGGAAC | CGTATTCACCTAGTTCTCTGCC |
| *NAMPT* | AAGCTGTTCCTGAGGGCTTT | TGTGGCCACTGTGATTGGAT |
| *HMGA1* | GCAGCCTCCGGTGAGT | GCACCCTTGTTTTTGCTTCC |
| Unmethylated *BRCA1*-qMSP | TTGGTTTTTGTGGTAATGGAAAAGTGT | CAAAAAATCTCAACAAACTCACACCA |
| Methylated *BRCA1*-qMSP | AAAACGACAATACAAAAACCGTC | TAGAGATAGCGGTAGAGTTGGTAGC |
| Bisulfite *BRCA1* | ATTTTTGTGGGGTGAATTTAATATG | CCCTCAACCCCAATATTTATTATTT |
| *RhoA* | CGTTAGTCCACGGTCTGGTC | ACCAGTTTCTTCCGGATGGC |
| *ActinB* | AGCACAGAGCCTCGCCTTT | GAAGCCGGCCTTGCACAT |
| **Taqman assays** | | |
| **Gene** | **Taqman Primer catalog #** | **Company** |
| *BRCA1* | Hs01556193_m1 | Thermo Fisher Scientific, Pleasanton, CA, USA |
| *PPIA* | Hs04194521_s1 | Thermo Fisher Scientific, Pleasanton, CA, USA |
| **Antibodies** | | |
| **Protein** | **Catalog No. and dilution** | **Company** |
| BRCA1 | Sc-6954, 1:1000, RRID:AB_626761 | Santa Cruz Biotechnology, Inc., Oregon, USA |
| ALDH1 | 611194, 1:1000, RRID:AB_2224312 | BD Transduction Laboratories, USA |
| Phospho-Histone H2A.X | 9718, 1:1000, RRID:AB_2118009 | Cell Signaling Technology, Danvers, MA, USA |
| ActinB | 8457, 1:2000, RRID:AB_10950489 | Cell Signaling Technology, Danvers, MA, USA |
| DNMT1 | D4692, 1:1000, RRID:AB_262096 | Sigma- Aldrich, St. Louis, MO, USA |
| DNMT3B | HPA001595, 1:1000, RRID:AB_1847814 | Sigma- Aldrich, St. Louis, MO, USA |
| Phospho-Histone H3 | 9701, 1:50, RRID:AB_331535 | Cell Signaling Technology, Danvers, MA, USA |
| Phospho-ATM | 13050, 1:1000, RRID:AB_2798100 | Cell Signaling Technology, Danvers, MA, USA |

### Supplementary Methods

#### Generation of cisplatin resistant OVCAR5 cells

To generate cisplatin resistant cells, OVCAR5 cells were treated with IC70 dose of cisplatin (18 μM) for 3 hours. Following treatment, media was removed, cells were washed two times with PBS, fresh media without cisplatin was added back to the flask and the cells were allowed to recover. Once cells reached 60-70% confluency, cells were split at 1:2. When cells reached 80% confluency the cisplatin treatment was repeated as the second cycle of treatment. Cells received 5 cycles of treatment and a maintenance treatment with 18 μM cisplatin was performed every month to retain a resistant population.

#### Mitotic Index analysis

After treatment, cells were trypsinized and resuspended in 1X cold PBS. Cells were fixed by drop-wise addition of absolute ethanol while vortexing and stored at -20 °C for 24 h or until all samples were collected. After washing cells with 1X PBS, phospho-Histone H3 (Ser10) antibody (1:50 dilution, 9701, Cell Signaling, RRID:AB_331535) was added and solution was rotated for 1 h at 37 °C in antibody dilution reagent (5% BSA in 1X PBS). Then, after washing cells with antibody reagent, fluorochrome conjugated secondary antibody AF-488 (1:1000 dilution, 4412, Cell Signaling, RRID:AB_1904025) was added and solution was incubated for 30 min at 37 °C in dark. Next, cells were washed with cold 0.1% BSA in 1X PBS and permeabilized in cold 0.1% Triton X in 1X PBS. PI stain (11348639001, Roche Diagnostics, Indianapolis, USA) was added in 100 ⌠g/mL RNAase A and incubated for 30 min at 37°C in dark. After subsequent washing with 1X PBS, cells were resuspended and filtered through a 30 mm filter (Sysmex).

Flow cytometry analysis was performed on a LSRII flow cytometer (BD Biosciences) at IU Flow Cytometry Core Facility. PI and pH3 stain were measured using 488 nm excitation and the signal was detected using the 610/20 and 530/30 filter respectively and analyzed at least three times in independent experiments. Additionally, samples with no stain and PI only and pH3-AF488 only labeling were analyzed for every experiment for compensation control. For each experiment, 10,000 events were analyzed. Further sample analysis was done in FlowJo software (Becton, Dickinson & Company).
